# Dorsal skull meningeal lymphatic vessels drain blood-solutes after intracerebral hemorrhage

**DOI:** 10.1101/2021.03.09.434530

**Authors:** Anaïs Virenque, Raz Balin, Francesco M. Noe

## Abstract

Drainage of intraparenchymal hematoma is crucial for the treatment of intracerebral hemorrhage (ICH). We investigated here the possible function of the meningeal lymphatic vessels (mLVs) in ICH resolution. Meningeal lymphatics have been reported to be involved in cerebrospinal fluid drainage, but their role in the drainage and clearance of brain parenchyma has not been characterized in details. Using two preclinical models of ICH, mimicking focal cortical ischemic hemorrhage and subcortical extended hemorrhage, we characterized the dynamics of blood drainage through the dorsal mLVs by two-photon real time imaging in awake mice. After ICH induction, we observe the flow of blood-derived components within the mLVs, suggesting that meningeal lymphatics can play a role in facilitating the drainage of the hemorrhage. We also found that local formation of new mLVs is directly correlated with ICH-related neuroinflammation levels. These findings suggest that meningeal lymphatics could provide a valuable therapeutic target for ICH resolution.

**Summary:** In vivo awake imaging reveals the direct drainage of extravasated blood-solutes from brain parenchyma into dorsal meningeal lymphatic vessels, following focal or diffuse intracranial hemorrhage

## Introduction

Intracerebral hemorrhage (ICH) occurs due to a cerebral vessel breach, causing blood leakage into the brain parenchyma (Schrag and Kirshner, 2020). It represents the most common form of brain hemorrhage (Kathirvelu et al., 2015) and the subtype of stroke with the highest mortality rate (Thabet et al., 2017). ICH is a severe, often lethal, neurological condition leading to chronic disabilities (Li et al., 2020): hematoma expansion and related secondary edema are associated with neurological deterioration and poor clinical outcomes, indicating a main therapeutic target in the control of the hematoma (Thabet et al., 2017). Currently, approved treatments for ICH are aimed at decreasing the hematoma mass effect, by optimizing blood pressure, and/or correcting the coagulopathy, with surgical intervention being the primary emergency choice. However, surgical intervention is not always possible (due to cause, location and size of the ICH), and has shown only minimal effects in neurological recovery (Chen et al., 2015; Schrag and Kirshner, 2020). Over the past few decades, many therapeutic approaches have been investigated, but few of them have been shown to be clinically relevant for the treatment of ICH (Napier et al., 2019; Zhao et al., 2020; Chen et al., 2015), and no clinically-approved drug treatments exist so far.

A functional system of lymphatic vessels has been recently described in the meninges surrounding the brain and the spinal cord (Aspelund et al., 2015; Louveau et al., 2015). Experimental evidences have indicated that the meningeal lymphatic vessels (mLVs) are part of the drainage routes for CNS fluids (cerebrospinal fluid – CSF, and interstitial fluid – ISF), and for the brain clearance (Sun et al., 2018; Louveau et al., 2016).

We propose that mLVs can be involved in the drainage of blood after cortical and subcortical hemorrhage: our hypothesis is that mLVs are recruited after ICH to contribute to the drainage of the blood from the brain parenchyma, constituting an important exit route that can alleviate hematoma growth and accelerate its resolution. To test our hypothesis, we used two *in vivo* models of ICH in mice: (1) intracerebral injection of collagenase IV, resulting in a subcortical diffused hemorrhage (Masuda et al., 2010; Rosenberg et al., 1990); (2) a laser-induced disruption of blood vessel wall (targeted photo-disruption), mimicking a focal cortical ischemic hemorrhage (Allegra Mascaro et al., 2010). ICH models have been induced in Prox1-eGFP mice that express GFP in lymphatic endothelial cells (Antila et al., 2017). Blood flow and ICH generation have been imaged by labeling the blood vessels with a vital dye (70 KDa tetramethylrhodamine-dextran; TRITC-dextran). Using two-photon microscopy, we characterized in real time *in vivo* the dorsal mLV routes and the dynamics of blood-derived tracer outflow, before and during the 120 min following the ICH induction (**Fig. 1**).

**Fig 1:**
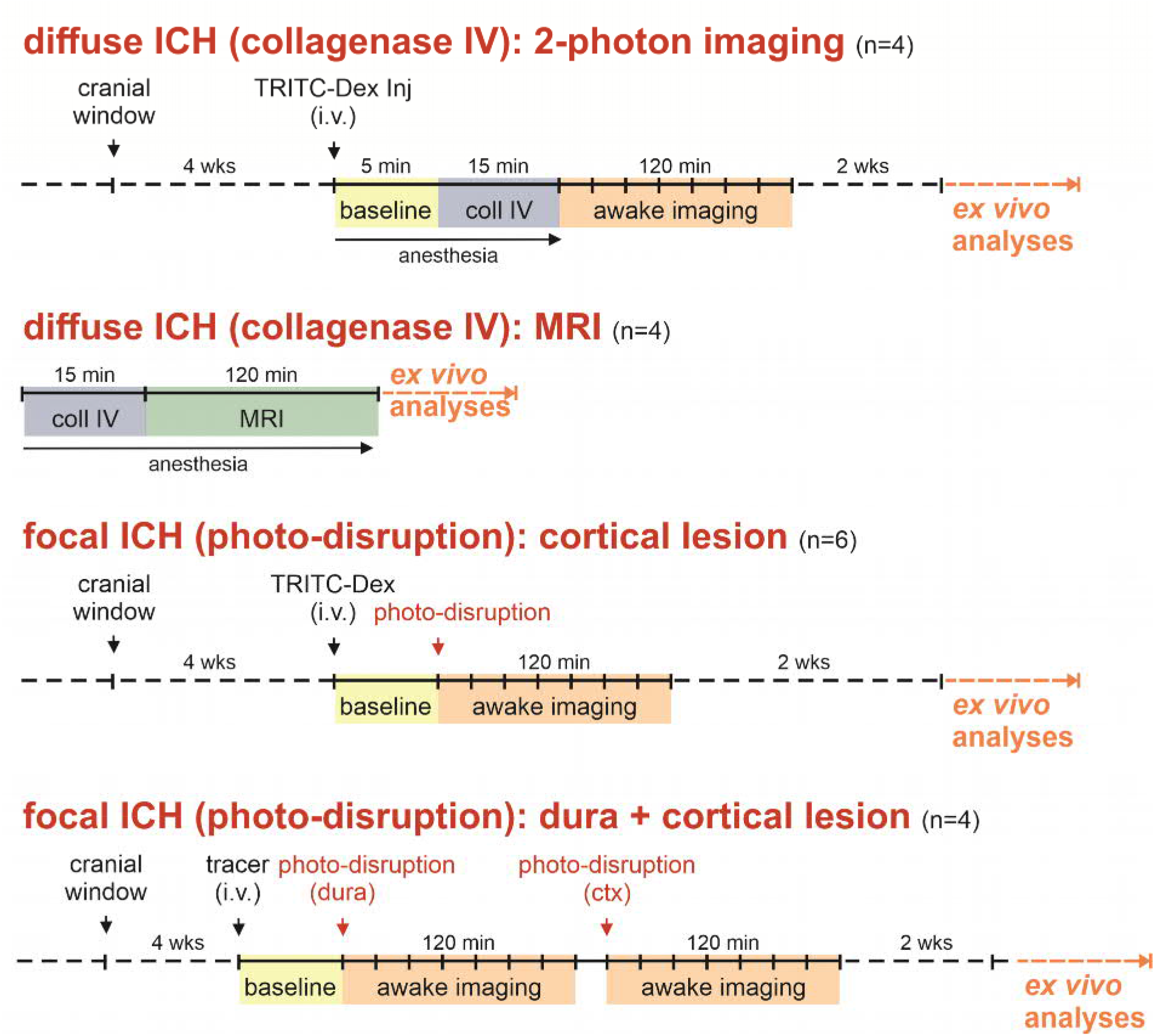
Study design. Schematic representation of the different experimental approaches used in the study. Number (n) of animals used in each experiment is reported within brackets. In the diffuse ICH (collagenase IV) experiments, animals were anesthetized with ketamine-medetomidine for the retro-orbital i.v. tracer injection, and during the intracortical collagenase IV administration. For mLV 2-photon imaging, mice were awakened by the administration of atipamezole. For MRI acquisition, mice were anesthetized for the whole duration of the study. During 2-photon imaging, eight images have been acquired for each location, every 15 min. For methodological details, refer to the Methods section.

## Results

### Blood-solutes drain through mLVs after diffuse ICH

To assess the involvement of mLVs in ICH resolution, we first analyzed the blood drainage through mLVs after the induction of an extensive subcortical hemorrhage. To mimic this condition, we injected collagenase IV into the superior colliculus (**Fig. 2A**). Within 10 min from injection, intracranial bleeding was visible across the whole superior colliculus and extended towards the retrosplenial granular cortex (**Fig. 2B**). Blood vessels, hemorrhage formation and blood-solutes drainage in the mLVs were imaged in awake mice, using TRITC-dextran as reporter tracer. After ICH induction, continuous imaging of Prox1 positive mLV (located around the confluence of sinuses) showed a significant increase in TRITC-dextran signal inside the lymphatic lumen starting from 37.5 min (± 12.0 min) after collagenase IV injection, and a peak at 60 min (p = 0.035 uptake *vs*. baseline; linear mixed-effect model with Bonferroni correction) (**Fig. 2C, D and F**). As reported before, intracerebral bleeding persists for hours after collagenase IV administration(Kirkman et al., 2011): at the latest time point analyzed in our experimental design (120 min after ICH induction) we still observed the presence of a strong tracer signal inside the imaged mLV (**Fig. 2E**). However, we also observed an increase of the tracer signal in the underlying parenchyma, resulting in a decrease of the calculated ∆P intensity (p = 0.036 uptake *vs*. 120 min; linear mixed-effect model with Bonferroni correction) (**Fig. 2E and F**). Our data clearly show the involvement of dorsal mLVs in the drainage of blood solutes after an extended subcortical hemorrhage. On the other hand, collagenase IV model of ICH is associated with acute edema formation, changes in intracranial pressure, accumulation of erythrocytes in the perivascular space and in the white matter fiber tracts, and tissue necrosis (Rosenberg et al., 1990). Moreover, type IV collagen is found in the basal lamina surrounding the brain blood vessels, as well as in the Virchow-Robin space (Rosenberg et al., 1990). It is therefore possible that the injection of collagenase IV results in a modification of the physiological flow dynamics of the ISF, eventually resulting in a passive flow of blood solutes in the dorsal mLVs.

**Fig 2:**
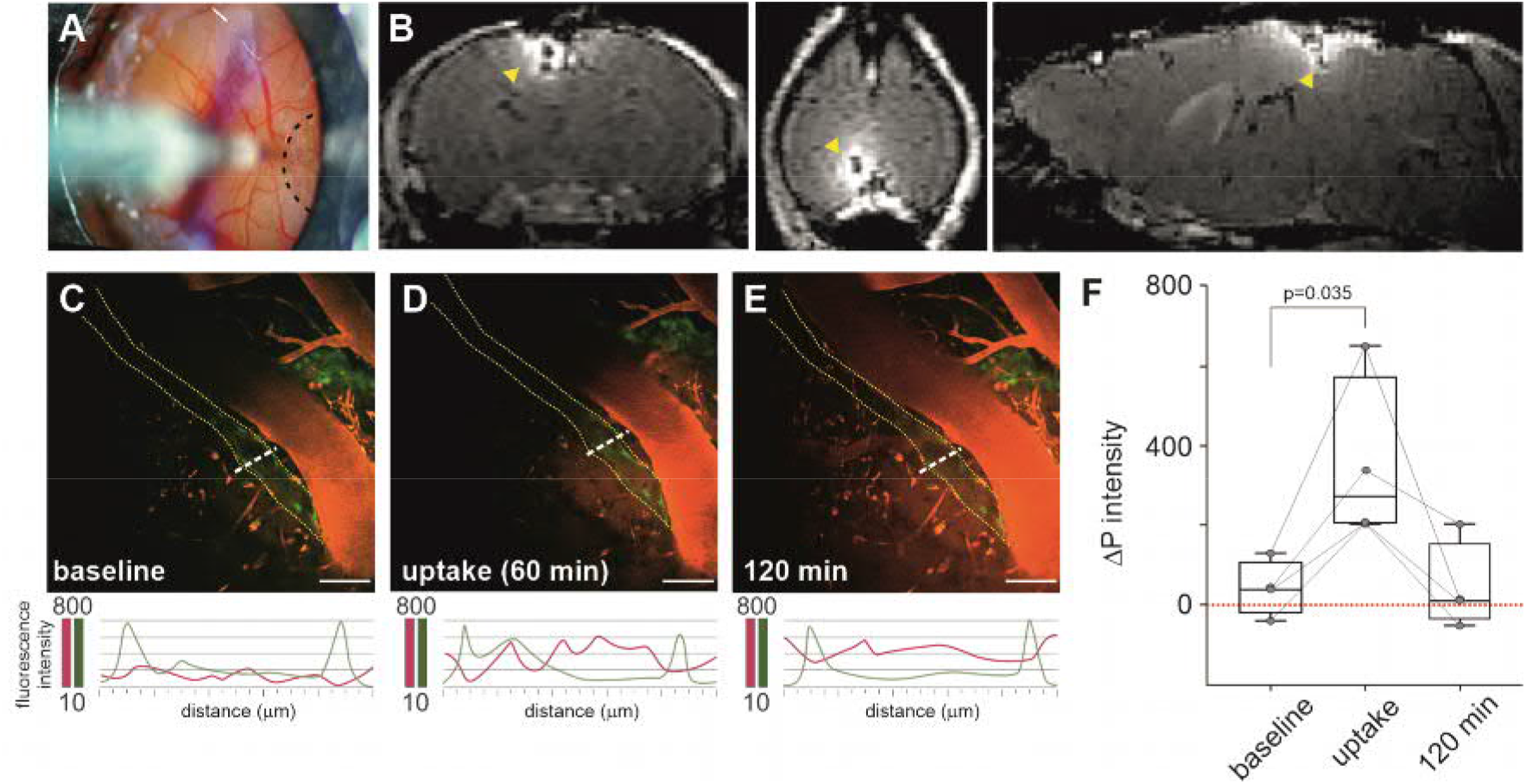
Blood-derived solutes drain through the mLV after severe ICH in deep cortical and subcortical layers, induced by collagenase IV injection. **(A)** In Prox1-eGFP reporter mice, collagenase IV is injected with a glass capillary in the deep cortical layer (2 mm underneath dura), through a pokable cranial window (black dotted line). **(B)** Representative coronal (left), axial (middle) and sagittal (right) reconstructed T2-weighted MRI images showing the hemorrhage extension in the deep cortical layers (yellow arrowhead), 120 min after collagenase IV injection (n = 4). **(C-E)** *Top panels:* Representative 2-photon imaging from the dura mater (second harmonic generation – blue) of TRITC-dextran containing blood vessels (red), and of a likely collector mLV (green). Images have been acquired during the same recording session, before collagenase injection (baseline; **C**), and 60 min (uptake; **D**) or 120 min (**E**) following ICH induction (n = 4). The dotted yellow line defines the profile of the imaged mLV. Panels **D** and **E** show the presence of TRITC-positive signal in the mLV lumen, suggesting that the mLVs are actively involved in the drainage of blood solutes from the parenchyma, after a severe ICH. *Bottom panels:* Intensity profiles of the TRITC tracer signal centered on the mLVs, as analyzed along the dashed white line in **C, D** and **E** respectively, showing the peak intensities for each channel (green: eGFP expressing lymphatic endothelial cells; red: TRITC-dextran tracer). Note that, after ICH induction, the TRITC-dextran signal peaks between the two eGFP-peaks (which define the mLV lumen), indicating the uptake of the leaked tracer by the mLV. **(F)** ΔP indicates the difference between the TRITC-dextran intensities as measured within and outside the mLV. Analysis of ΔP variations at different time points before and after ICH induction shows a statistically significant uptake of the leaked TRITC-dextran tracer by the mLVs. Apparent return to baseline levels, as measured at 120 min after ICH induction, is due to the increase of tracer signal in the underlying parenchyma, resulting from the sustained intracerebral hemorrhage. Gray lines connect values from the same mouse. Scale bar in C-E: 30µm. Data analysis: linear mixed-effect model with Bonferroni correction.

### mLVs drain blood-solutes from a focal cortical ICH, but not from a subdural hemorrhage

Next, we analyzed if dorsal mLVs are engaged in blood drainage also after a focal, transient cortical hemorrhage. Focal ICH was induced in awake mice by a targeted photo-disruption of a superficial cortical blood vessel using high-intensity pulsed laser, generated by the 2-photon microscope (Allegra Mascaro et al., 2010). The rupture of the targeted small vessel resulted in an immediate increase of the TRITC-dextran fluorescent signal in the brain parenchyma, confirming the induction of ICH. Imaging of the tracer signal within the mLVs ipsilateral to the lesion, showed a signal increase in the lymphatic lumen after 8.0 minutes (± 4.5 min), with maximum intensity observed at 65 min after photo-disruption (**Fig. 3A-D; Movie S1**). The signal intensity decreased over time and was completely absent 120 min after the transient small vessel rupture (p = 0.015 uptake *vs*. baseline; p = 0.015 uptake *vs*. resolution by linear mixed-effect model with Bonferroni correction) (**Fig. 3D**). Analysis of the TRITC-dextran signal within the mLV lumen, imaged along the lymphatic antero-posterior length, showed a gradual and unidirectional filling of the vessel (**Fig. 3E-H**): tracer filled the mLV downstream to the site of hemorrhage, apparently toward the major lymphatics located around the transverse sinuses and the cerebellar ring (Louveau et al., 2018b). Time-lapse imaging of mLV during the uptake phase further confirmed the unidirectional lymphatic flow towards the caudal portion of the analyzed mLVs (**Movie S2**).

**Fig 3:**
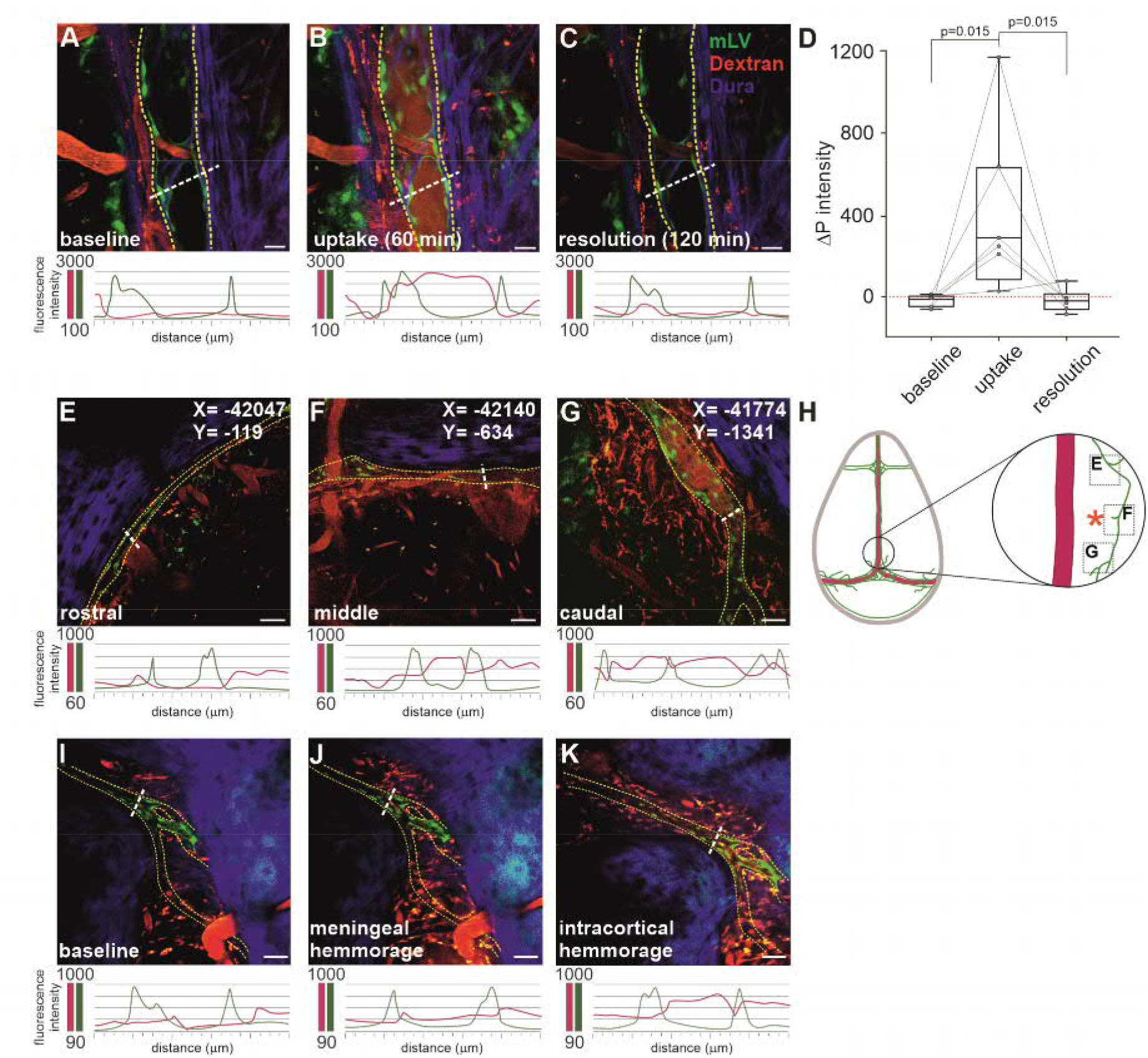
Selective mLV drainage after focal cortical but not meningeal ICH. **(A-C)** Representative 2-photon imaging of tracer-containing blood vessels (red) and mLV (green), in the dura mater (second harmonic generation – blue), at baseline condition **(A)**, after focal cortical ICH **(B)** and after hemorrhage resolution **(C)**, and respective intensity profiles (green: eGFP expressing lymphatic endothelial cells; red: TRITC-dextran tracer) (n = 6). At baseline, no TRITC-dextran signal is detected within the mLV. After ICH induction, local meningeal lymphatics uptake and drain the blood solute from the brain parenchyma, as inferable by the presence of tracer signal within the lumen of the mLV. After hemorrhage resolution, no signal is evident within the mLV, suggesting an active transport of blood solutes through the lymphatics. Scale bars: 15 µm. **(D)** Analysis of the difference between the TRITC-dextran intensities (as measured within and outside the mLV; ΔP) shows a significant increase of the tracer signal in the mLV after ICH, compared to the baseline. After ICH resolution the ΔP reverts to baseline levels. **(E-G)** Representative 2-photon images, and respective intensity plots, of an mLV along its antero-posterior axis, collected 60 min after focal ICH induction. No tracer signal within the mLV is visible in the rostral portion of the vessel, upstream to the site of lesion **(E)**. Tracer signal in the mLV lumen (indicating blood-solutes uptake) is observed in the middle **(F)** and caudal **(G)** sections of the lymphatic, suggesting a unidirectional flow directionality. Absolute coordinates of the imaged areas are reported in each panel. Scale bars: 50 µm. **(H)** Schematic representation of the mLV (green), running in proximity of the sinus sagittalis (red), imaged in **E-G**. Red asterisk indicate the location of the induced focal ICH. **(I-K)** mLV imaged at baseline **(I)**, after dural blood vessel lesion **(J)** or after focal ICH **(K)** (n = 4). As visible in panel **J**, mLV do not uptake the tracer after a localized meningeal hemorrhage. However when the hemorrhage is induced in the underlying cortex, a clear TRITC-dextran signal is visible within the lymphatic lumen, suggesting that mLVs take-up and drain blood-solutes from the brain parenchyma and not from the dura. Scale bars: 40 µm. Yellow dotted line defines the profile of the imaged mLVs. Data analysis: linear mixed-effect model with Bonferroni correction.

Our data indicate that, independent of the ICH etiology, mLVs located in the confluence of sinuses region are directly involved in the blood clearance from the brain parenchyma. However, we could not assume that there was no spillover of the tracer into the subarachnoid space after ICH induction. This consideration is important as recent works (Pu et al., 2019; Chen et al., 2020) have suggested the involvement of mLVs in the drainage of blood after subarachnoid hemorrhage (Chen et al., 2020). To investigate the possible tracer uptake directly from the meningeal space, we induced a focal transient hemorrhage in the dura, by targeting a dural blood vessel located in proximity of the imaged mLVs (**Fig. 3H-I**). No increase of TRITC-dextran signal was observed in the lymphatic lumen during the 120 min of imaging (p = 0.250 uptake *vs*. baseline by Wilcoxon matched-pairs signed rank test), suggesting that mLVs do not actively participate in the blood drainage from the meningeal space (**Fig. 3I**). In order to confirm the functionality of the imaged mLV and its ability to uptake blood solutes, 150 min after the induction of the dural hemorrhage, in the same mouse we induced a focal IHC. This resulted in tracer uptake inside the mLV lumen, with a kinetic comparable to the one described above (**Fig. 3J**). These results suggest that, in our experimental conditions, the dorsal mLVs are involved in the blood drainage directly and specifically from the parenchyma.

### ICH-induced neuroinflammation correlates with focal mLVs lymphangiogenesis

Formation of new lymphatic vessels (lymphangiogenesis) has been reported after implantation of an epidural electrode or a cranial window (Hauglund NL, Kusk P, Kornum BR, Nedergaard M, 2020). In our experiments, the cranial window implantation *per se* did not induce lymphangiogenesis, as longitudinally evaluated *in vivo* at 2 and 4 weeks after the window implantation (**Fig. 4A and B**). Data were confirmed by *ex vivo* analysis of whole-mounted meninges, performed 6 weeks after cranial window surgery (**Fig. 4C**). At the same time, no neuroinflammation in the underlying brain tissue was observed associated with the implant of the cranial window (**Fig. 4G**), excluding the hypothesis that our experimental preparation could result in the alteration of physiological conditions (*i*.*e*., glymphatic flow)(Hauglund NL, Kusk P, Kornum BR, Nedergaard M, 2020). Similarly, the analysis of tissues from focal ICH animals, collected 2 weeks after injury, did not reveal lymphangiogenesis in the dura mater (**Fig. 4D**), and only a minor activation of astrocytes and microglia could be observed in some animals, confined to the cortical areas adjacent to the ICH site (**Fig. 4H**). In contrast, diffuse ICH resulted in an increase of Prox1 positive vessels around the site of collagenase IV injection (p = 0.013 *vs*. control by Kruskall-Wallis test) (**Fig. 4E and F**), in association with a pronounced reactive gliosis (microglia: p = 0.005 *vs*. control; astrocytes: p = 0.021 *vs*. control by Kruskall-Wallis test) (**Fig. 4I**). We also found a direct correlation between the lymphatic coverage and neuroinflammation levels (**Fig. 4L and M**), suggesting that sustained, but not transient, inflammatory events in the brain modulate the lymphatic vessels in the meninges.

**Fig 4:**
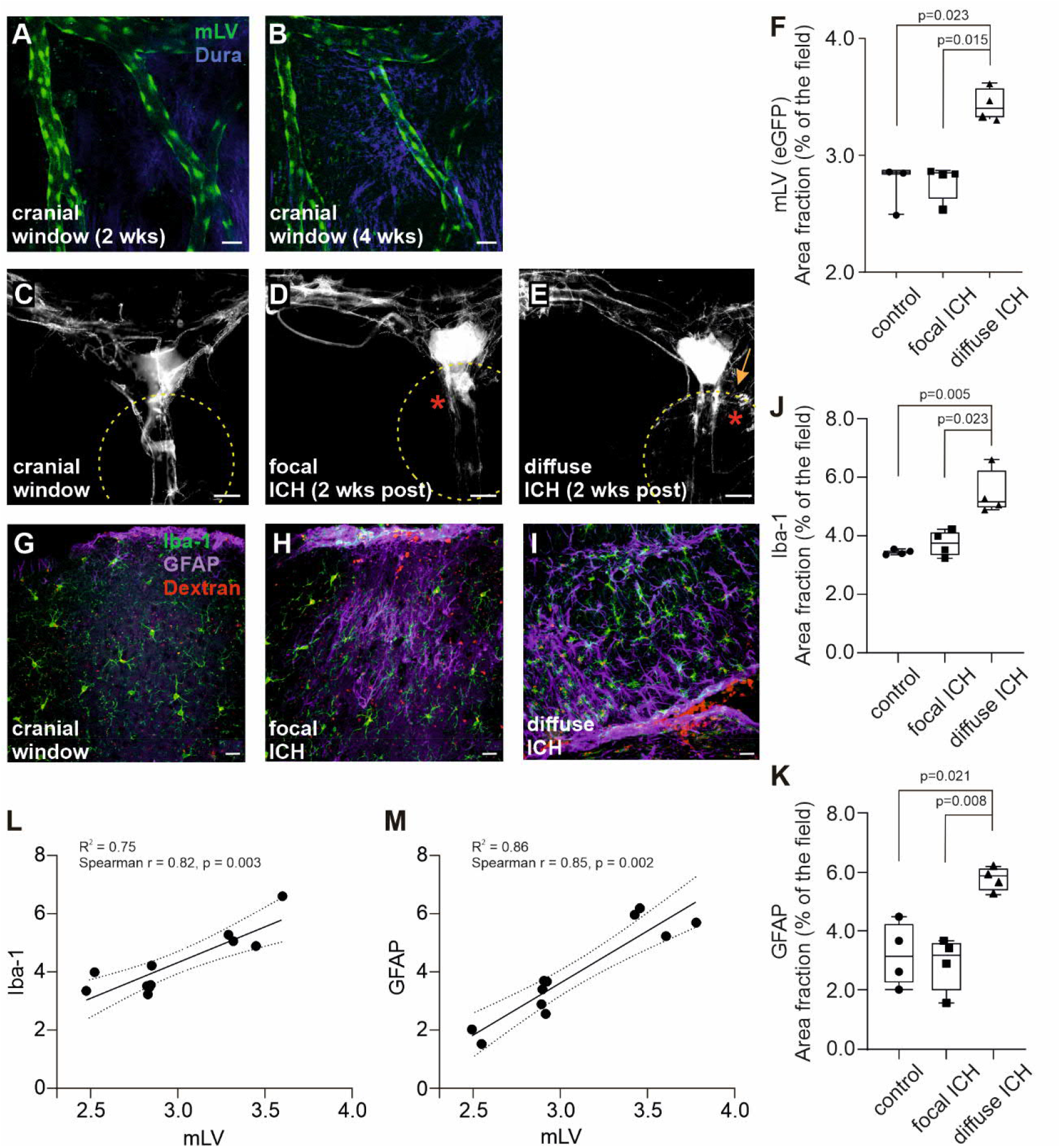
Chronic cranial window, as well as focal or diffuse ICH do not affect mLV lymphangiogenesis, despite being associated with different neuroinflammation levels. **(A-B)** Representative in vivo 2-photon imaging of Prox1-eGFP-positive mLV (green), 2 weeks (**A**) and 4 weeks (**B**) after cranial window implantation. No mLV sprouting or changes in mLV conformation were observed related to the implant of the cranial window. Scale bar: 30 µm. **(C-E)** Representative images of meningeal whole mounts after cranial window implant or after ICH induction, from Prox1-eGFP mice. (**C**) 6 weeks after cranial window and no injury (cranial window; n = 4). (**D**) 2 weeks after photo-disruption (focal ICH; n = 4). (**E**) 2 weeks after collagenase IV injection (diffuse ICH; n = 4). Meningeal lymphangiogenesis has been observed after diffuse ICH (arrow in panel **E**). Scale bars: 50 µm. **(F)** Percent area of Prox1-eGFP coverage at 6 weeks post cranial window implant, or 2 weeks post focal or diffuse ICH (n = 4 each experimental group). **(G-I)** Representative confocal image of gliosis (microglia: Iba1, green; astrocytes: GFAP, violet) at the site of ICH, 2 weeks after lesion induction. No reactive gliosis was observed associated with the cranial window per se (**G**). Minor microglia and astrocyte activation could be observed after focal ICH (**H**), while diffuse ICH corresponded to extensive gliosis (**I**). Scale bars: 20 µm. **(J-M)** Quantification of the percent area of Iba-1 (**J**) and GFAP (**K**) immunoreactivity, and respective correlation analyses with the lymphatic coverage (percent area). (**L**) Iba-1, (**M**) GFAP. Neuroinflammation directly correlates with the increase in lymphatic coverage of the dorsal meninges. Data analyses: Kruskal-Wallis test with Bonferroni correction and Spearman correlation.

## Discussion

Here, using live imaging in awake mice, we analyzed the involvement of dorsal meningeal lymphatics during intracerebral hemorrhage. Our work supports the hypothesis that meningeal lymphatics are involved in the drainage (and possibly clearance) of blood from brain parenchyma after ICH (Da Mesquita et al., 2018; Aldea et al., 2016). We also expand our understanding on mLV function, providing evidences that the dorsal mLVs serve as a direct and specific conduit for the transport of solutes from the brain parenchyma. Finally, we report a correlation between the levels of neuroinflammation and meningeal lymphangiogenesis, contributing to demonstrate that glia activation modulates the meningeal lymphatics. The role of dorsal meningeal lymphatics as a drainage pathway for the CNS has been disputed, with most of the studies providing indirect evidences of the transport through mLVs, and/or focusing on the cerebro-spinal fluid (CSF) clearance (for a review see (Proulx, 2021)). In controlled and clinical-mimicking conditions, we found a unidirectional transport of blood-derived solutes through mLVs, which suggests a direct flow from the brain parenchyma to the mLV. At the same time, our data excludes the involvement of dorsal mLVs in the immediate drainage of blood solutes from the meningeal space, therefore revising the current paradigm that meningeal lymphatics are important for the CSF drainage.

Our data apparently challenge a recent publication, which proposed a role for dorsal mLVs in the resolution of subarachnoid hemorrhage (Chen et al., 2020). Chen and colleagues reported the presence of erythrocytes in the dorsal lymphatics starting from 4 h after the injection of 60 μl of autologous blood into the cisterna magna. This time-course, however, is unlikely compatible with the possibility that mLVs directly uptake the erythrocytes from the CSF. Rather, it suggests that the recirculation of CSF (Kress et al., 2014) and a secondary uptake from the perivascular and/or the Virchow Robin space are involved.

A functional coupling between the mLVs and the so-called glymphatic system has been previously proposed (Louveau et al., 2017; Rasmussen et al., 2018; Sun et al., 2018; Da Mesquita et al., 2018). Our data support this hypothesis and contribute to demonstrate that this coupling is important for the drainage of blood after ICH. After collagenase IV injection, despite a more extended and severe hemorrhage, we observed a clear delay in the tracer uptake than after focal photo-disruption. Different factors can prompt the observed delay: intrinsic features of the collagenase IV ICH model (such as edema formation, increase of intracranial pressure, perivascular space obstruction and pia degradation) have been shown to affect the glymphatic flow. Moreover, the induction of this model requires the use of anesthesia (**Fig. 1**), which has been reported to modulate the glymphatic system (Hablitz et al., 2019). All these factors can impair the glymphatic flow, affecting the drainage of blood solutes in the parenchyma and ultimately delaying the tracer uptake by the mLVs.

Finally, our analyses revealed a direct correlation between the level of neuroinflammation and the extension of lymphatic coverage in the corresponding area of dorsal meninges. These data expand and consolidate previous indirect reports (Hauglund NL, Kusk P, Kornum BR, Nedergaard M, 2020; Louveau et al., 2018b). We speculate that local meningeal lymphatic plasticity, in response to neuroinflammatory state of the underlying brain, can contribute to regulate fluid homeostasis and neuro-immune surveillance in the brain. However, further studies are needed to characterize the mechanisms of this interaction, and the role of the increased lymphangiogenesis in brain allostasis, which may be relevant in different neurological conditions.

This is a limited, descriptive study, conducted on two clinically relevant animal models, revealing the dorsal meningeal lymphatics as a specific route for the drainage of blood-solutes after ICH. Yet, the characterization of the lymphatic flow presented here serves as a template for future explorations of mLV drainage, with promising implications for ICH treatment. Drug treatments (such as administration of VEGF-c)(Song et al., 2020), or other strategies aimed at increasing the meningeal lymphatic flow, could be evaluated in experimental and clinical conditions, possibly representing innovative and effective therapies for the treatment of acute ICH and associated neuroinflammation. Overall, our work expands the knowledge on the function of meningeal lymphatics and provides a model for studying the relation between mLVs and the glymphatic system, as well as their contribution in different neurological conditions.

## Materials and Methods

### Study design

This was an *in vivo* descriptive study, conducted in awake mice, designed to characterize the drainage and flow of blood-solutes after focal or diffuse ICH. Study design, including experimental groups and number of individual biological replicates are summarized in **Fig. 1**. No data, including outlier values, were excluded. Sample sizes used for each experiment are detailed in the figure legends. Investigators were not blinded for any part of the study.

### Animals

All the experiments were approved by the Animal Experiment Board of the State Provincial Office of Southern Finland, Finland (ESAVI/008787/2017; ESAVI/13987/2018), and performed in accordance with the University of Helsinki animal care regulations.

Five months old male and female Prox1-eGFP mice (Antila et al., 2017) (backcrossed more than 10 times to C57BL/6JOlaHsd background mice) have been used in this study. A total of 22 mice have been used: n = 6 for targeted photo-disruption ICH; n = 4 for meningeal hemorrhage + targeted photo-disruption ICH; n = 4 for collagenase IV-induced ICH; n = 4 for MRI assessment of ICH after collagenase IV administration; and n = 4 for assessment of lymphangiogenesis related to cranial window implant.

### Cranial window implant

Mice were anesthetized using a fresh prepared solution of Domitor® (medetomidine -1.0 mg/kg) -Ketaminol® (ketamine -75 mg/kg) in sterile saline by i.p. injection. During surgery, mice were placed on a heating blanket connected to a temperature controller, to maintain the animal core temperature at 35.6°C. Dexamethasone (2 mg/kg, s.c.) was administered the day before surgery, as well as soon before the start of surgery procedures and 24 h after, to prevent inflammation. Carprofen (4 mg/Kg, s.c.) was administered before surgery as painkiller. In order to protect eyes from dehydration and irritation, ophthalmic ointment was applied. Fur on the surgery region was shaved, scalp scrubbed three times with 70% ethanol and 10% betadine (povidone-iodine), and Xylocain gel was applied locally. After making a cutaneous midline incision over the skull and removing the periosteum, skull was degreased with acetone and rinsed with sterile saline. Craniotomy was manually performed with a 3 mm trephine: craniotomy was centered over the midline suture, with the posterior edge apposed over the lambda, in front of pineal gland (**Fig. 2A and 4C**). In order to reduce heating during craniotomy, the skull was irrigated with cold artificial CSF (aCSF). The craniotomy was filled with sterile aCSF and covered with a circular coverglass (3 mm diameter, #1 thickness; IS-Vet), which was glued to the skull with cyanoacrylate glue (Loctite 401; Henkel). Centered on the top of the glass, a stainless steel head plate (Pryazhnikov et al., 2018) was fixed using glue and epoxy cement (total C-RAM; ITENA), paying attention not to damage the imaging window. The head plate is essential to stabilize the animal under the two-photon microscope for the subsequent imaging sessions. At the completion of surgical procedure, the animal was injected s.c. with the specific medetomidine-antagonist Atipamezole (Antisedan® vet 5mg/mL) at the dose of 1 mg/kg. Pre-warmed sterile saline (1.0 mL i.p., at 32°C) was administered to favor post-anesthesia recovery, and the animal was placed in an incubator at 32.5°C for 1 h, before being transferred to its own homecage. Transparency of the cranial window was assessed every week starting from the second week post-surgery.

### Two-photon imaging

Mice were imaged using a FV1200MPE two-photon microscope (Olympus, Japan), equipped with a 25X water immersion 1.05 NA objective specially designed for *in vivo* two-photon imaging. MaiTai Broad Band DeepSee laser (tuned to 880 nm) was used for excitation. Emission light has been collected through a band pass filter (515–560 nm). For *in vivo* imaging sessions, awake animals were head-fixed under the 2-photon microscope using a Mobile HomeCage device (Neurotar). Mobile HomeCage allows the movement of the animal during the imaging session and at the same time minimizes movement-related artefacts, facilitating stable imaging without the use of anesthesia. Prior to the imaging sessions, animals have been habituated to head fixation in the mobile HomeCage, during eight 2 h-long training sessions (Pryazhnikov et al., 2018). Three-dimensional (3D) baseline GFP fluorescence image Z-stacks of the mLVs were acquired. Z-stacks of images have been collected with the vertical step size of 2 µm (total depth of imaged tissue: 50–100 µm) with zoom factor 1 at 800 × 800 pixels aspect ratio. For each animal, up to 3 different mLVs were selected, imaging 3 regions of interest (ROIs) for each vessel along its antero-posterior route.

### Targeted photo-disruption of blood vessel wall

Four weeks after cranial window implant, baseline images of mLVs were acquired in n = 10 mice (n = 6 cortical photo-disruption; n = 4 dura + cortical photo-disruption) after retro-orbital injection of 100 μl TRITC-dextran to visualize the blood vasculature. After acquiring images at resting state, a blood capillary located in the superficial cortical layer (± 100 μm below the bottom margin of the dura) or in the inferior layer of the dura (localized using the second harmonic generation), and within 100 μm from the imaged mLV, were selected. The spatial localization provided by multi-photon excitation was used to target specifically the blood vessel, being careful that during photo-disruption the laser did not cross the mLVs. Photo-disruption was achieved by focusing the laser beam at the blood-vessel wall to induce a highly precise and reproducible local injury. Using the Olympus Fluoview software, a linear ROI was drawn across the wall of the targeted vessel and the insult was generated by irradiation of the vessel with a train of amplified 1-kHz, 100-fs, 800 nm laser pulses(Nishimura et al., 2006) set at 100% of energy. Irradiation time (*i*.*e*., number of pulses) was increased proportionally during subsequent sessions until the haemorrhage was observed. We started below the threshold of observed disruption to the vessel: if no vascular changes were observed, we increased the laser power and repeated irradiation. Mean duration of irradiation to achieve haemorrhage in the cortex was 3 ± 2 sec. After photo-disruption, ICH formation was briefly imaged for 10 sec, to confirm blood vessel rupture. This protocol induces a focal transient hemorrhage, as observed in our experimental conditions, where tracer leakage from the damaged vessels subsided within 10 min from the photo-disruption induction. Imaging plane was then moved to the dura to image mLVs, as previously described. Z-stack images of each ROI were acquired every 5 min. To visualize the flow within the mLVs, time-lapse images were acquired at 10 sec intervals. A total of 40 images were acquired (Rosen et al., 2001; Allegra Mascaro et al., 2010).

### Intracortical collagenase injection

Mice used for multiphoton imaging (n = 4) experiments were implanted with a special 5 mm cranial window equipped with a silicone access port (Roome and Kuhn, 2014). Four weeks after cranial window implant, baseline z-stack images of mLVs were acquired following retro-orbital administration of 100 μl TRITC-dextran. Thereafter, mice were anesthetized using a freshly prepared solution of Domitor®-Ketaminol®, and positioned on a stereotaxic frame for collagenase IV injection: 0.5 μL of collagenase IV (0.15U/μL solution in aCSF; Gibco) was injected in the superior colliculus, through the silicone access port (Roome and Kuhn, 2014) (**Fig. 2A**). Collagenase IV was delivered through a 50 μm bore-silicate capillary connected to a 10 μL Hamilton syringe, at the constant rate of 0.1 μL/min using a programmable micropump (Legato200, KD scientific). At the completion of collagenase IV delivery, the injection capillary was left in place for an additional 10 min for proper diffusion of the solution and to avoid back-flow. Thereafter, animals were awakened by s.c. administration of Antisedan®, and immediately connected to the 2-photon microscope for imaging (total anesthesia duration: 20 min).

A different cohort of mice (n = 4) were used to image collagenase IV-related ICH by MRI. Briefly, the animals were anesthetized by Domitor® -Ketaminol®, fixed in a stereotaxic frame, and the skull has been exposed. 200 μl of Gadovist® solution was injected i.v. (retro-orbital administration). A burr hole was drilled at the coordinates (referred to bregma): -2,8 mm anteroposterior, and +1 mm mediolateral. A 50 μm bore-silicate capillary filled with collagenase IV (0.15U/μL solution in aCSF) was lowered through the hole to -2 mm from the dura surface (dorsoventral coordinate). Collagenase IV was injected as described before (0.5 μL at 0.1 μL/min), and the animal was placed on an MR-compatible holder with head fixed to minimize movement artefacts during scanning. MRI data have been collected at 7T (Bruker Pharmascan, Ettlingen, Germany) using 3D T2-weighted Fast Spin-Echo sequence (RARE, repetition time 1.5 s, effective echo time 48 ms, 16 echoes per excitation) and 100 μm isotropic resolution (field of view 25.6 mm x 128.8 mm x 9.6 mm). Acquisition matrix was 128 x 256 x 96. Throughout the imaging session (approx. 2 h), the mice have been kept under anesthesia. Body core temperature was maintained at 35°C. A pressure sensor was used to monitor the respiratory rate, and respiratory-gating was used to minimize motion artifacts. At the end of MRI acquisition, mice were euthanized by intracardial perfusion with cold PBS followed by 4% paraformaldehyde (PFA), and brains and meninges collected for further analyses.

### Sample preparation for histological analyses

Two weeks after ICH induction, mice were intracardially perfused with ice-cold PBS (6 ml/min; 6 min) followed by 4% PFA (6 ml/min, 10 min). In order to collect meninges for whole-mount preparation and brain for histological analyses, skulls were harvested as previously described (Antila et al., 2017). Dorsal and basal skullcaps, and brain were post-fixed in 4% PFA for 12 h. Thereafter, brains were cryoprotected in 30% sucrose for 48 h, frozen with *N*-pentane on dry ice, embedded into OCT compound, and stored at –80 °C. 25 μm coronal sections were cut by cryostat, and collected in solution containing 30 % ethylene glycol, 25 % glycerol in 0.05 M phosphate buffer (PB) and stored at -20 °C until further processing. Fixed meninges (dura mater and arachnoid) were carefully dissected from the dorsal skullcaps with forceps and kept in PBS at 4 °C until further use (Louveau et al., 2018a).

### Immunohistochemistry and imaging

#### Image acquisition from Prox1-eGFP meninges

Image acquisition was performed using a Zeiss Axio Observer Z1 microscope, equipped with a Zeiss AxioCam MR R3 camera, mounting a 10x lens to obtain images from whole-brain sections (Antila et al., 2017).

*GFAP, Iba1 and DAPI*. For immunofluorescence procedure, brain sections were washed and blocked in a blocking solution (4 % BSA, 0,2 % Triton X-100 in PBS) for 1 h at RT, followed by overnight incubation at 4 °C with the following primary antibodies diluted in blocking solution: mouse anti-GFAP (1:500, Sigma G3893), and rabbit anti-Iba1 (1:500, Wako 01-1974). After washing in PBS, sections were incubated for 2 h at RT with secondary fluorescent antibodies in blocking solution: Alexa Fluor 546-conjugated goat anti mouse (1:250), Alexa Fluor 647-conjugated donkey anti rabbit (1:250 all from Invitrogen, Thermo Fisher Scientific). Thereafter, sections were washed in PBS before being mounted onto glass slides and cover slipped using Vectashield® with DAPI (BioNordika Oy). High-resolution Z-stack images were captured with a confocal microscope (Zeiss LSM710), using 25X objective. ZEN 2012 software (Carl Zeiss GmbH) was used for image processing.

### Image analysis

After acquisition, the multiphoton images were processed and analyzed using Imaris (Bitplane) and Fiji/ImageJ (open source image processing package, NIH, USA) software. For the uptake after ICH, the intensity of Prox1-eGFP and rhodamine dextran signal was measured using the profile plot analysis of Imaris software. A line was drawn through the lymphangion segment to demonstrate the change in intensity of the rhodamine-dextran signal over time in the lumen of the mLV (Fig.1). The Δ*P* intensity was calculated as the difference between the peak value of rhodamine-dextran signal inside the mLV and the rhodamine-dextran signal outside the mLV (Ahn et al., 2019).

For immunochemistry analysis, two representative brain sections from the site of the lesion (approximately –2.18 to -3.16 bregma) or the corresponding area in sham animals were imaged. The contour of the section was traced as region of interest (ROI), and the threshold was uniformly set for each experiment to select for stained cells (Iba1+ and GFAP+ cells). For assessment of meningeal lymphatic vessel coverage, the entire meningeal whole-mount preparation was threshold and used for quantification. The percent area coverage of Prox1 was used to determine the coverage of the lymphatic vessels. The mean percent area fraction was calculated using Microsoft Excel (Bolte et al., 2020).

### Statistical analysis

Statistical analyses were performed using R v3.5.3 software/computing environment (The R foundation for statistical computing). All software packages were taken from the Comprehensive R Archive Network mirror sites (CRAN; http://CRAN.R-project.org/package=boot). A linear mixed model with restricted maximum likelihood (lmer in lme4 package) was used for the analyses of ΔP intensity. Graphs have been designed using GraphPad Prism v8.4.2 (graphPad softwares). Bonferroni correction was used to adjust p-values in multiple comparison. Significance was accepted at the level of p < 0.05. Data are expressed as median ± SD.

## Acknowledgments

For their help with MRI sequences, the authors would like to thank Mikko Kettunen and Riikka Immonen from Biomedical Imaging Unit, National Bio-NMR facility, A.I.Virtanen Institute for Molecular Sciences, University of Eastern Finland. Authors thank also Marja Lohela (BioImaging Unit, Biomedicum, University of Helsinki), Leonard Khirug, Marina Tibeykina and Evgeny Pryazhnikov (Awake Rodent Research Facility – ARRP, HiLIFE-Neuroscience Center, University of Helsinki) for their help and training with two-photon microscope. Authors thank Prof. Mikko Airavaara for his help with collagenase IV protocol, and Prof. Kari Alitalo and Dr. Salli Antila for their contribution in discussing the conclusions of this work.

## Author contributions

AV: Conceptualization, Methodology, Investigation, Validation, Formal analysis, Data Curation, Writing – Review and Editing; RB: Investigation, Validation; FMN: Conceptualization, Methodology, Validation, Writing, Supervision, Funding acquisition.

## Data and materials availability

The raw data supporting the conclusions of this manuscript will be made available by the corresponding author, upon reasonable request, to any qualified researcher.

## Supplementary Materials

**Movie S1**. 3D reconstruction of tracer uptake in the mLV lumen after focal ICH (10 min).

**Movie S2**. mLV flow after focal ICH (45 min-time lapse).

### Supplementary Video Captions

**Movie S1: Imaris three-dimensional (3D) reconstruction of 2-photon microscopy imaging**. To visualize blood-solutes uptake by mLVs, TRITC-dextran was injected i.v. before focal ICH induction. The 3D reconstruction of a 2-photon Z-stack image, acquired 10 min after focal ICH induction in an awake freely-moving mouse, shows a specific signal from TRITC-dextran tracer (red) in the mLV lumen (green).

**Movie S2: in vivo 2-photon imaging time-lapse showing the lymphatic drainage of TRITC-dextran tracer (red) and of putative immune cells in the mLV lumen (green)**. Putative immune cells can be observed as black dots moving within the mLV lumen (green), following the direction of the lymphatic flow. Representative time-lapse video of 3 independent experiments: photograms have been taken every 10 sec.

## References

Ahn, J.H., H. Cho, J.H. Kim, S.H. Kim, J.S. Ham, I. Park, S.H. Suh, S.P. Hong, J.H. Song, Y.K. Hong, Y. Jeong, S.H. Park, and G.Y. Koh. 2019. Meningeal lymphatic vessels at the skull base drain cerebrospinal fluid. Nature. 572:62–66. doi:10.1038/s41586-019-1419-5.

Aldea, R., B. Bedussi, A.W.J. Morris, R.O. Weller, and R.O. Carare. 2016. Lymphatic Clearance of the Brain?: Perivascular, Paravascular and Significance for Neurodegenerative Diseases. Cell. Mol. Neurobiol. 36:181–194. doi:10.1007/s10571-015-0273-8.

Allegra Mascaro, A.L., L. Sacconi, and F.S. Pavone. 2010. Multi-photon nanosurgery in live brain. Front. Neuroenergetics. 2:1–8. doi:10.3389/fnene.2010.00021.

Antila, S., S. Karaman, H. Nurmi, M. Airavaara, M.H. Voutilainen, T. Mathivet, D. Chilov, Z. Li, T. Koppinen, J.-H. Park, S. Fang, A. Aspelund, M. Saarma, A. Eichmann, J.-L. Thomas, and K. Alitalo. 2017. Development and plasticity of meningeal lymphatic vessels. J. Exp. Med. 214:3645–3667. doi:10.1084/jem.20170391.

Aspelund, A., S. Antila, S.T. Proulx, T.V. Karlsen, S. Karaman, M. Detmar, H. Wiig, and K. Alitalo. 2015. A dural lymphatic vascular system that drains brain interstitial fluid and macromolecules. J. Exp. Med. 212:991–999. doi:10.1084/jem.20142290.

Bolte, A.C., A.B. Dutta, M.E. Hurt, I. Smirnov, M.A. Kovacs, C.A. McKee, H.E. Ennerfelt, D. Shapiro, B.H. Nguyen, E.L. Frost, C.R. Lammert, J. Kipnis, and J.R. Lukens. 2020. Meningeal lymphatic dysfunction exacerbates traumatic brain injury pathogenesis. Nat. Commun. doi:10.1038/s41467-020-18113-4.

Chen, J., L. Wang, H. Xu, L. Xing, Z. Zhuang, Y. Zheng, X. Li, C. Wang, S. Chen, Z. Guo, Q. Liang, and Y. Wang. 2020. Meningeal lymphatics clear erythrocytes that arise from subarachnoid hemorrhage. Nat. Commun. 11:1–15. doi:10.1038/s41467-020-16851-z.

Chen, S., Q. Yang, G. Chen, and J.H. Zhang. 2015. An Update on Inflammation in the Acute Phase of Intracerebral Hemorrhage. Transl. Stroke Res. 6:4–8. doi:10.1007/s12975-014-0384-4.

Hablitz, L.M., H.S. Vinitsky, Q. Sun, F.F. Stæger, B. Sigurdsson, K.N. Mortensen, T.O. Lilius, and M. Nedergaard. 2019. Increased glymphatic influx is correlated with high EEG delta power and low heart rate in mice under anesthesia. Sci. Adv. doi:10.1126/sciadv.aav5447.

Hauglund NL, Kusk P, Kornum BR, Nedergaard M. 2020. Meningeal lymphangiogenesis and enhanced glymphatic activity in mice with chronically implanted EEG electrodes. J Neurosci. doi:10.1523/JNEUROSCI.2223-19.2020.

Kathirvelu, B., S.T. Carmichael, and L. Angeles. 2015. Intracerebral hemorrhage in mouse models: interventions and functional recovery. Metab. Brain Dis. 30:449–459. doi:10.1007/s11011-014-9559-7.Intracerebral.

Kirkman, M.A., S.M. Allan, and A.R. Parry-Jones. 2011. Experimental intracerebral hemorrhage: avoiding pitfalls in translational research. J. Cereb. Blood Flow Metab. 31:2135–2151. doi:10.1038/jcbfm.2011.124.

Kress, B.T., J.J. Iliff, M. Xia, M. Wang, and H. Wei. 2014. Impairment of paravascular clearance pathways in the aging brain. Ann Neurol. 76:845–861. doi:10.1002/ana.24271.

Li, L., R. Luengo-Fernandez, S.M. Zuurbier, N.C. Beddows, P. Lavallee, L.E. Silver, W. Kuker, and P.M. Rothwell. 2020. Ten-year risks of recurrent stroke, disability, dementia and cost in relation to site of primary intracerebral haemorrhage: Population-based study. J. Neurol. Neurosurg. Psychiatry. 91:580–585. doi:10.1136/jnnp-2019-322663.

Louveau, A., A.J. Filiano, and J. Kipnis. 2018a. Meningeal whole mount preparation and characterization of neural cells by flow cytometry. Curr Protoc Immunol. 121:1–10. doi:10.1002/cpim.50.

Louveau, A., T.H. Harris, and J. Kipnis. 2015. Revisiting the concept of CNS immune privilege. Trends Immunol. 36:569–577. doi:10.1016/j.it.2015.08.006.Revisiting.

Louveau, A., J. Herz, M.N. Alme, A.F. Salvador, M.Q. Dong, K.E. Viar, S.G. Herod, J. Knopp, J.C. Setliff, A.L. Lupi, S. Da Mesquita, E.L. Frost, A. Gaultier, T.H. Harris, R. Cao, S. Hu, J.R. Lukens, I. Smirnov, C.C. Overall, G. Oliver, and J. Kipnis. 2018b. CNS lymphatic drainage and neuroinflammation are regulated by meningeal lymphatic vasculature. Nat. Neurosci. 21. doi:10.1038/s41593-018-0227-9.

Louveau, A., B.A. Plog, S. Antila, K. Alitalo, M. Nedergaard, and J. Kipnis. 2017. Understanding the functions and relationships of the glymphatic system and meningeal lymphatics. J. Clin. Invest. 127:3210–3219. doi:10.1172/JCI90603.

Louveau, A., I. Smirnov, T.J. Keyes, J.D. Eccles, J. Sherin, J.D. Peske, N.C. Derecki, D. Castle, J.W. Mandell, L. Kevin, T.H. Harris, and J. Kipnis. 2016. Structural and functional features of cns lymphatics. Nature. 523:337–341. doi:10.1038/nature14432.Structural.

Masuda, T., M. Maki, K. Hara, T. Yasuhara, N. Matsukawa, S. Yu, E. Cate, N. Tajiri, S.H. Chheda, M. Aurora, N. Weinbren, Y. Kaneko, S.A. Kirov, D.C. Hess, H. Hida, and C. V Borlongan. 2010. Perihemorrhagic degeneration accompanies stereotaxic collagenase-mediated cortical hemorrhage in mouse. Brain Res. 1355:228–239. doi:10.1016/j.brainres.2010.07.101.

Da Mesquita, S., Z. Fu, and J. Kipnis. 2018. The Meningeal Lymphatic System: A New Player in Neurophysiology. Neuron. 100:375–388. doi:10.1016/j.neuron.2018.09.022.

Napier, J., L. Rose, O. Adeoye, E. Hooker, K.B. Walsh, J. Napier, L. Rose, O. Adeoye, E. Hooker, and B. Kyle. 2019. Modulating acute neuroinflammation in intracerebral hemorrhage?: the potential promise of currently approved medications for multiple sclerosis promise of currently approved medications for multiple sclerosis. Immunopharmacol. Immunotoxicol. 41:7–15. doi:10.1080/08923973.2019.1566361.

Nishimura, N., C.B. Schaffer, B. Friedman, P.S. Tsai, P.D. Lyden, and D. Kleinfeld. 2006. Targeted insult to subsurface cortical blood vessels using ultrashort laser pulses: Three models of stroke. Nat. Methods. 3:99–108. doi:10.1038/nmeth844.

Proulx, S.T. 2021. Cerebrospinal fluid outflow: a review of the historical and contemporary evidence for arachnoid villi, perineural routes, and dural lymphatics. Cell. Mol. Life Sci. 1:3. doi:10.1007/s00018-020-03706-5.

Pryazhnikov, E., E. Mugantseva, P. Casarotto, J. Kolikova, S.M. Fred, D. Toptunov, R. Afzalov, P. Hotulainen, V. Voikar, R. Terry-Lorenzo, S. Engel, S. Kirov, E. Castren, and L. Khiroug. 2018. Longitudinal two-photon imaging in somatosensory cortex of behaving mice reveals dendritic spine formation enhancement by subchronic administration of low-dose ketamine. Sci. Rep. doi:10.1038/s41598-018-24933-8.

Pu, T., W. Zou, W. Feng, Y. Zhang, L. Wang, H. Wang, and M. Xiao. 2019. Persistent malfunction of glymphatic and meningeal lymphatic drainage in a mouse model of subarachnoid hemorrhage. Exp. Neurobiol. 28:104–118. doi:10.5607/en.2019.28.1.104.

Rasmussen, M.K., H. Mestre, M. Nedergaard, and M. Sciences. 2018. The glymphatic pathway in neurological disorders. Lancet Neurol. 17:1016–1024. doi:10.1016/S1474-4422(18)30318-1.

Roome, C.J., and B. Kuhn. 2014. Chronic cranial window with access port for repeated cellular manipulations, drug application, and electrophysiology. Front. Cell. Neurosci. 8:1–8. doi:10.3389/fncel.2014.00379.

Rosen, E.D., S. Raymond, A. Zollman, F. Noria, M. Sandoval-Cooper, A. Shulman, J.L. Merz, and F.J. Castellino. 2001. Laser-induced noninvasive vascular injury models in mice generate platelet-and coagulationdependent thrombi. Am. J. Pathol. 158:1613–1622. doi:10.1016/S0002-9440(10)64117-X.

Rosenberg, G.A., S. Mun-bryce, M. Wesley, and M. Kornfeld. 1990. Collagenase-Induced Intracerebral Hemorrhage in Rats. Stroke. 04:801–807. doi:10.1161/01.str.21.5.801.

Schrag, M., and H. Kirshner. 2020. Management of Intracerebral Hemorrhage: JACC Focus Seminar. J Am Coll Cardiol. 75:1819–1831. doi:10.1016/j.jacc.2019.10.066.

Song, E., T. Mao, H. Dong, L.S.B. Boisserand, S. Antila, M. Bosenberg, K. Alitalo, J.L. Thomas, and A. Iwasaki. 2020. VEGF-C-driven lymphatic drainage enables immunosurveillance of brain tumours. Nature. 577:689–694. doi:10.1038/s41586-019-1912-x.

Sun, B.L., L. hua Wang, T. Yang, J. yi Sun, L. lei Mao, M. feng Yang, H. Yuan, R.A. Colvin, and X. yi Yang. 2018. Lymphatic drainage system of the brain: A novel target for intervention of neurological diseases. Prog. Neurobiol. 163–164:118–143. doi:10.1016/j.pneurobio.2017.08.007.

Thabet, A.M., M. Kottapally, and J.C. Hemphill. 2017. Management of intracerebral hemorrhage. Handb. Clin. Neurol. 140:177–194. doi:10.1016/B978-0-444-63600-3.00011-8.

Zhao, W., C. Wu, C. Stone, Y. Ding, and X. Ji. 2020. Treatment of intracerebral hemorrhage?: Current approaches and future directions. J. Neurol. Sci. 416:117020. doi:10.1016/j.jns.2020.117020.

